# Structural changes in the SARS-CoV-2 spike E406W mutant escaping a clinical monoclonal antibody cocktail

**DOI:** 10.1101/2022.01.21.477288

**Authors:** Amin Addetia, Young-Jun Park, Tyler Starr, Allison J. Greaney, Kaitlin R Sprouse, John E. Bowen, Sasha W. Tiles, Wesley C. Van Voorhis, Jesse D. Bloom, Davide Corti, Alexandra C. Walls, David Veesler

## Abstract

The SARS-CoV-2 receptor-binding domain (RBD) E406W mutation abrogates neutralization mediated by the REGEN-CoV therapeutic monoclonal antibody (mAb) COVID-19 cocktail and the cilgavimab (AZD1061) mAb. Here, we show that this residue substitution remodels the ACE2-binding site allosterically, thereby dampening receptor recognition severely and altering the epitopes recognized by these three mAbs. Although vaccine-elicited neutralizing antibody titers are decreased similarly against the E406 mutant and the Delta or Epsilon variants, broadly neutralizing sarbecovirus mAbs, including a clinical mAb, inhibit the E406W spike mutant.

## Main

The receptor-binding domain (RBD) of the severe acute respiratory syndrome coronavirus 2 (SARS-CoV-2) spike glycoprotein is responsible for interacting with the host receptor ACE2 and initiating viral entry into cells^1–3^. The SARS-CoV-2 RBD is the target of the majority of neutralizing antibodies elicited by SARS-CoV-2 infection and COVID-19 vaccination as well as monoclonal antibodies (mAbs) used therapeutically^4–8^. Binding and neutralization of SARS-CoV-2 by individual mAbs can be escaped by single RBD residue mutations, which led to the development of therapeutic cocktails comprising two mAbs recognizing non-overlapping epitopes^9–12^. These cocktails have a higher barrier for the emergence of neutralization escape mutants than the individual constituting mAbs, as typically at least two distinct amino-acid substitutions are required to evade neutralization by a two-mAb cocktail.

The REGEN-COV cocktail consists of two mAbs, casirivimab (REGN10933) and imdevimab (REGN10987) that bind non-overlapping RBD epitopes in the receptor-binding motif (RBM), and block ACE2 attachment^11,12^. We previously mapped all possible RBD residue mutations that permit escape from the REGEN-COV mAb cocktail and the cilgavimab (AZD1061) mAb which led us to identify that the E406W substitution abrogated binding and neutralization of both REGEN-COV mAbs and the cocktail^9^ as well as binding of cilgavimab^13^. Unexpectedly, residue E406 is located outside of the epitopes recognized by casirivamab, imdevimab and cilgavimab, suggesting this mutation might influence the overall structure of the RBD (presumably through an allosteric effect) while retaining detectable binding to dimeric human ACE2^9^.

To understand the molecular basis of the E406W-mediated escape from the REGEN-COV cocktail and cilgavimab, we characterized the SARS-CoV-2 spike ectodomain trimer structure harboring the E406W mutation using single-particle cryo-electron microscopy. 3D classification of the dataset revealed the presence of two conformational states: one with three RBDs closed and one with one RBD open accounting for approximately 70% and 30% of particles, respectively. We determined a structure of the closed S state at 2.3 Å resolution applying C3 symmetry **(Figure 1, Figure S1 and Table 1)**. Symmetry expansion, focused classification and local refinement yielded an RBD reconstruction at 3.4Å resolution which was used for model building and analysis **(Figure 1, Figure S1 and Table 1)**.

**Figure 1.**
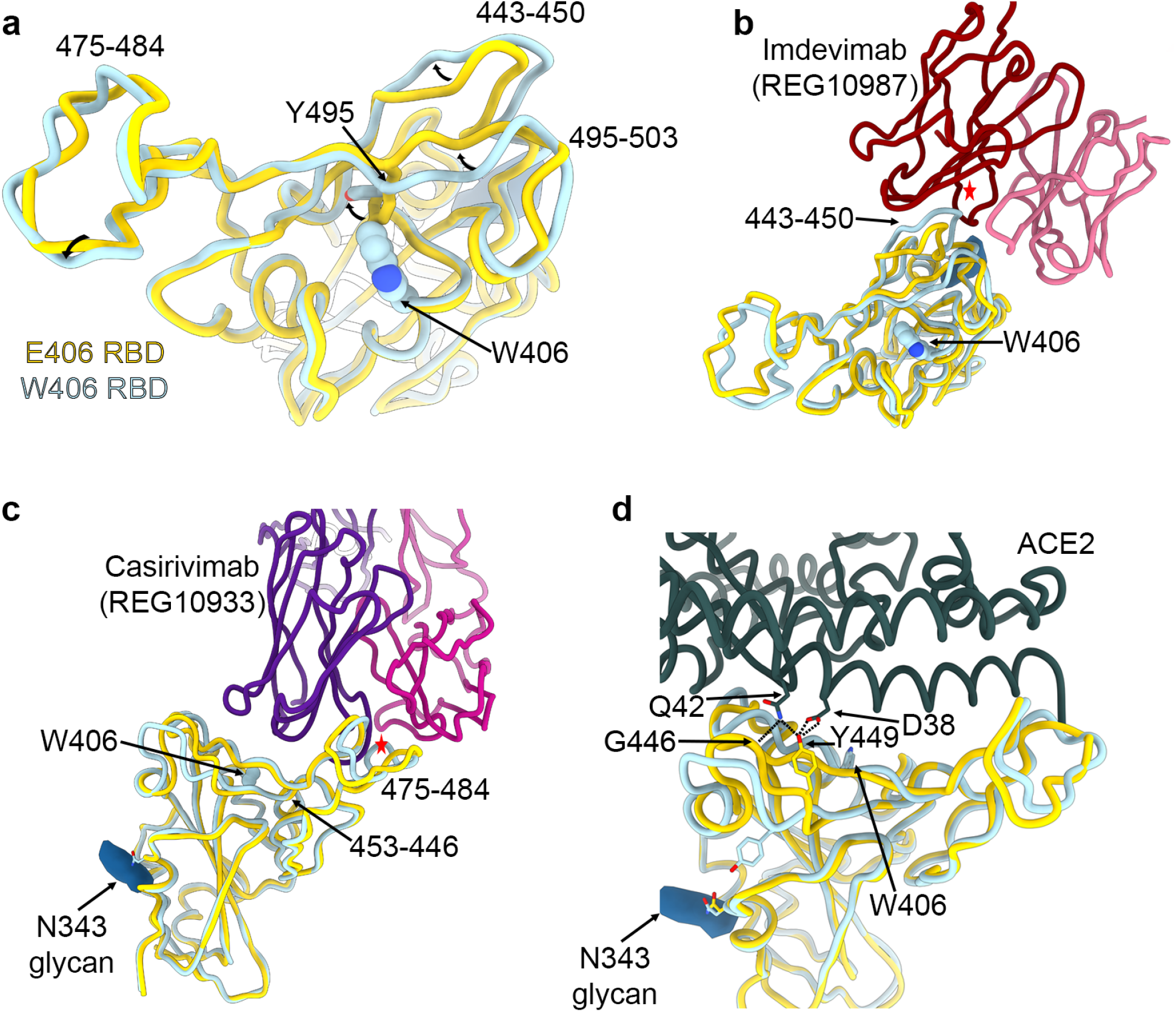
The E406W mutation remodels the SARS-CoV-2 RBD allosterically. **a**, Structural superimposition of the Wuhan-Hu-1 RBD (E406, gold, PDB 6m0j, ACE2 not displayed) and the W406 RBD (light blue). **b-c**, Structural superimposition of the imdevimab/casirivimab-bound Wuhan-Hu-1 RBD (E406, gold, PDB 6xdj) and the W406 RBD (light blue). Steric clashes indicated with red stars. **d**, Structural superimposition of the ACE2-bound Wuhan-Hu-1 RBD (E406, gold, PDB 6m0j) and the W406 RBD (light blue).Hydrogen bonds shown as dotted lines.

The E406W substitution places the introduced side chain indol ring in a position sterically incompatible with the neighboring Y495 phenol side chain, inducing a rotameric rearrangement of the latter residue relative to the ACE2-bound RBD structure^14^ or apo S ectodomain trimer structures^1,15^. This results in major conformational reorganization of residues 443-450 and 495-503 which experience up to 4.5Å shift relative to previously determined structures^1,15^. Although the organization of residues 475-484 are only subtly different in the E406W RBD relative to apo S structures^1,15^, it deviates markedly more from the ACE2-bound RBD structure^14^ or the REGEN-COV-bound RBD structure^11^ **(Figure 1a)**. Imdevimab (REGN10987) recognizes an epitope residing at the interface between antigenic sites Ia and IIa^5^ and forms extensive interactions with residues 440-449 that would sterically clash with the mAb heavy chain in the E406W RBD structure **(Figure 1b)**. Casirivamab (REGN10933) interacts with residues 417, 453-456 and 475-490 (within antigenic site Ia^5^) and the distinct conformation of the latter residues in the REGEN-COV-bound RBD and E406W apo S structures likely precludes mAb binding through steric clash with the mAb light chain **(Figure 1c)**. Our data therefore shows that the E406W mutation disrupts the antigenic sites recognized by casirivamab (REGN10933) and imdevimab (REGN10987) allosterically, which are positioned 5 and 20Å away, respectively^9^. Similar to imdevimab, the loss of cilgavimab (AZD1061) binding to the E406W RBD^13^ is explained by the structural reorganization of residues 443-450 which are recognized by this mAb **(Figure S2)**.

These RBD conformational changes also alter the ACE2-interacting surface resulting in the predicted loss of several hydrogen bonds formed between the ACE2 D38 and SARS-CoV-2 Y449 side chains as well as the ACE2 Q42 side chain and the SARS-CoV-2 Y449 side chain and G446 main chain carbonyl **(Figure 1d)**. Accordingly, we observed that the monomeric human ACE2 ectodomain bound with a 14-fold reduced affinity to immobilized SARS-CoV-2 E406W RBD (K_D_=1.34 µM) relative to wildtype (Wuhan-Hu-1) RBD (K_D_=93.9 nM) using biolayer interferometry **(Figure S3a-c and Table S2)**. This reduction of ACE2 binding affinity is expected to dampen viral fitness severely, as previously observed for another point mutation decreasing ACE2 binding^16^ **(Figure S3d)**.

A few broadly neutralizing sarbecovirus human mAbs have been recently described and shown to be resilient to the observed SARS-CoV-2 antigenic drift, to recognize distinct RBD antigenic sites, and protect small animals against challenge with SARS-CoV-2 variants of concern or other sarbecoviruses^10,16,18–22^. To evaluate the influence of the aforementioned structural changes on neutralization by these mAbs, we compared the concentration-dependent inhibition of S309, S2E12 and S2×259 against VSV particles pseudotyped with the G614 spike or the W406/G614 spike. Each of these three mAbs neutralized with comparable potency the G614 and W406/G614 pseudoviruses **(Table S3)**, indicating they retain activity against this mutant **(Figure 2A and Figure S4)**. As predicted based on structural data^5,10^, the S2H14 mAb failed to neutralize the spike W406/G614 pseudovirus due to the reorganization of the RBM **(Figure 2A and Figure S4)**. Moreover, these data are consistent with the fact that binding to the SARS-CoV-2 W406 RBD was unaffected for S2E12 and abrogated for S2H14^10^.

**Figure 2.**
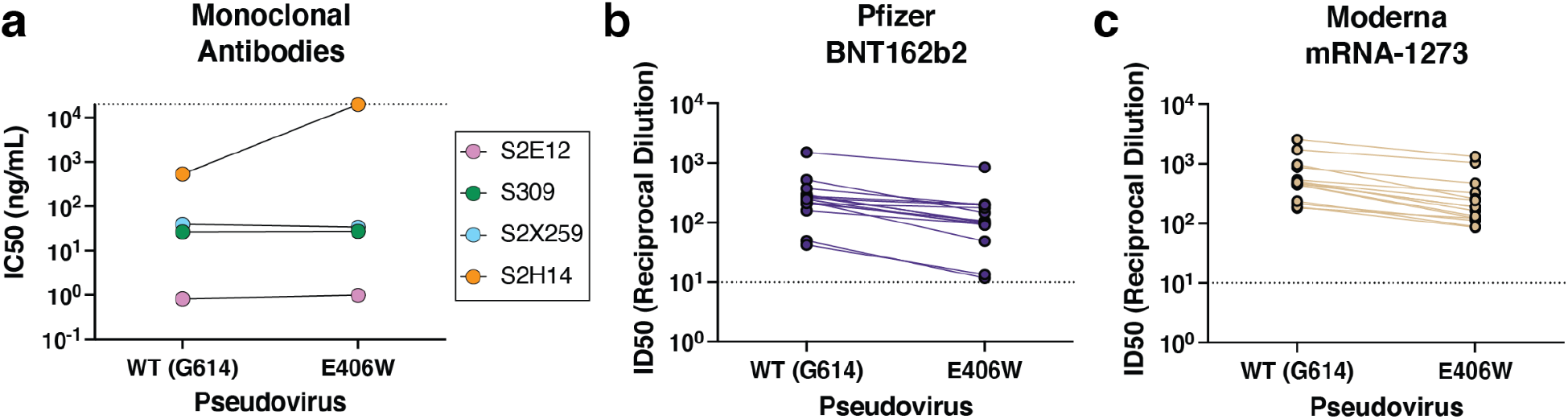
Evaluation of the neutralizing activity of several sarbecovirus broadly neutralizing mAbs and vaccine-elicited polyclonal antibodies. **a)** Neutralizationpotency potency (50% inhibition concentration, IC50) of the monoclonal antibodies S309, S2E12, S2×259, and S2H14 against VSV pseudotyped with either the wildtype (G614) or the E406W mutant spike protein. Non-neutralizing values are shown as 2 × 10^4^ ng/mL, the limit of detection of the assay, as indicated by a dotted line. **b-c)** Neutralization potency (50% inhibition dilution, ID50) of sera collected from individuals vaccinated with either Pfizer Cominarty (b) or Moderna’s mRNA-1273 (c) against VSV pseudotyped with SARS-CoV-2 wildtype (G614) or E406W spike. ID50 values measured against the two pseudoviruses for each sample are connected by a line. The dotted line indicates the limit of detection of the assay.

Finally, we set out to assess the impact of the E406W mutation on vaccine-elicited plasma neutralizing activity using samples obtained from individuals who had received 2 doses of either Pfizer BNT162b2 or Moderna mRNA-1273 COVID-19 vaccine **(Table S4)**. We observed 2.5-fold (BNT162b2, range: 1.2-4.6) and 2.4-fold (mRNA-1273, range: 1.5-3.8) reduction in neutralization potencies against the W406/G614 spike pseudovirus compared to G614 spike-harboring pseudovirus **(Figure 2B-C and Figure S5)**. These data indicate that the single E406W mutation leads to moderate erosion of vaccine-elicited polyclonal neutralizing antibodies, comparable to the SARS-CoV-2 Epsilon variant^23^ or the Delta variant^24^.

The ongoing SARS-CoV-2 genetic drift yielded variants harboring numerous mutations, some of them altering transmissibility, immune evasion, replication kinetics or disease severity relative to the ancestral SARS-CoV-2 isolate^7,23,25–307,23,25,30–37^. Although the E406W mutation promotes escape from REGEN-COV- and cilgavimab (AZD1061)-mediated neutralization, it requires multiple nucleotide substitutions from the Wuhan-Hu-1 spike sequence, has a strong deleterious effect on ACE2 binding and has not been detected in clinical isolates to date. Several best-in-class broadly neutralizing sarbecovirus mAbs are unaffected by the E406W mutation and COVID-19 mRNA vaccine-elicited polyclonal antibodies retain a substantial fraction of their activity against this mutant, indicating several strategies are available should a E406W mutant virus emerge in the future. Finally, our data showcase the structural and functional plasticity of the SARS-CoV-2 RBD^17^ that evolves under selective pressure from the host immune responses and the necessity to retain viral fitness of progeny viruses.

## Acknowledgements

We thank Hideki Tani (University of Toyama) for providing the reagents necessary for preparing VSV pseudotyped viruses. This study was supported by the National Institute of Allergy and Infectious Diseases (DP1AI158186 and HHSN272201700059C to DV, and R01AI141707 to JDB), the National Institute of General Medical Sciences (R01GM120553 to DV), the National Institute of Health Cellular and Molecular Biology Training Grant (T32GM007270 to A.A.), a Pew Biomedical Scholars Award (DV), an Investigators in the Pathogenesis of Infectious Disease Awards from the Burroughs Wellcome Fund (DV and JDB), Fast Grants (DV), the Bill & Melinda Gates Foundation (OPP1156262 to DV and INV-004949 to JDB), the University of Washington Arnold and Mabel Beckman cryoEM center and the National Institute of Health grant S10OD032290 (to D.V.) and grant U01 AI151698 for the United World Antiviral Research Network (UWARN) as part of the Centers for Research in Emerging Infectious Diseases (CREID) Network. TNS is an Howard Hughes Medical Institute Fellow of the Damon Runyon Cancer Research Foundation. JDB and DV are investigators of the Howard Hughes Medical Institute.

## Author contributions

Conceptualization: AA, T.N.S., A.J.G., J.B., A.C.W. and DV. Pseudovirus entry assays: AA and ACW. BLI measurements: AA. Provided unique reagents: D.C., SWT, WCVV Data analysis: AA and DV. Supervision: DV. Writing – original draft: AA and DV. Writing – review and editing: all authors

## Additional Information

Correspondence and requests for materials should be addressed to David Veesler.

## Methods

### Cell culture

Expi293 cells were grown in Expi293 media at 37°C and 8% CO_2_ rotating at 130 RPM. HEK-293T cells and HEK-293T cells stably expressing the human ACE2 receptor (HEK-ACE2)^38^ were grown in DMEM supplemented with 10% FBS and 1% PenStrep at 37°C and 5% CO_2_. Vero cells stably expressing the human protease TMPRSS2 (Vero-TMPRSS2) were grown in DMEM supplemented with 10% FBS, 1% PenStrep, and 8 µg/mL puromycin at 37°C and 5% CO_2_.

### Constructs

The construct encoding spike ectodomain harboring the E406W mutation was obtained from the Institute for Protein Design. The spike ectodomain was codon optimized, stabilized with the hexapro mutations^39^ and mutation of the furin cleavage site (_682_RRAR_685_ to _682_GSAS_685_), and inserted into the pCDNA3.1 vector containing a C-terminal foldon followed by an avi tag and an octa-histidine tag.

The construct encoding the E406W RBD was generated by performing around-the-horn mutagenesis using a pCMVR vector encoding the wildtype SARS-CoV-2 RBD containing an N-terminal mu-phosphatase signal peptide and a C-terminal avi tag and octa-histidine tag. The boundaries for the SARS-CoV-2 RBD in this construct were _328_RFPN_331_ to _528_KKST_531_.

### Recombinant protein expression and purification

To produce the SARS-CoV-2 spike ectodomain containing the E406W mutation, 125 mL of Expi293 cells were grown to density of 2.5 × 10^6^ cells per mL and transfected with 125 µg of DNA using PEI MAX diluted in Opti-MEM. The cells were grown for four days after which the supernatant was clarified by centrifugation. The recombinant ectodomain was purified using a nickel HisTrap FF affinity column, washed with 10 column volumes of 20 mM imidazole, 25 mM sodium phosphate pH 8.0, and 300 mM NaCl, and eluted with a 500 mM imidazole gradient. The purified proteins were buffer exchanged and concentrated in 20 mM sodium phosphate pH 8 and 100 mM NaCl using a 100 kDa centrifugal filter. The proteins were flash frozen and stored at -80°C until use.

The wildtype, B.1.1.7, and E406W RBDs were produced by transfecting 25 mL of Expi293 cells at a density of 2.5 × 10^6^ cells per mL with 25 µg of DNA using the ExpiFectamine 293 Transfection Kit. The cells were grown for four days and the resulting supernatant was collected and clarified by centrifugation. The recombinant RBD was purified using a nickel HisTrap HP affinity column, washed with 10 column volumes of 20 mM imidazole, 25 mM sodium phosphate pH 8.0, and 300 mM NaCl, and eluted using a 500 mM imidazole gradient. The resulting protein was buffer exchanged and concentrated using a 10 kDa centrifugal filter. Next, the purified RBDs were biotinylated using the BirA biotin-protein ligase reaction kit (Avidity). The biotinylated proteins were re-purified and concentrated as described above. The proteins were flash frozen and stored at -80°C until use.

### Cryo-EM sample preparation and data collection

Purified SARS-CoV-2 spike ectodomain harboring the E406W mutation was added to a freshly glow discharged 2.0/2.0 UltraFoil grid (200 mesh). The grid was then plunge frozen using a Vitrobot MarkIV (ThermoFisher) with a blotting force of 0 and time of 6.5 seconds at 100% humidity and 23°C. Data were acquired on a FEI Titan Krios transmission electron microscope operated at 300 kV and equipped with a Gatan K3 direct detector and Gatan Quantum GIF energy filter. Automated data acquisition was carried out using Leginon^40^. The dose rate was adjusted to 15 counts/pixel/s and each movie was acquired in 75 frames of 40 ms with pixel size of 0.843 Å and a defocus range comprised between 0 and -2.6 µm.

### CryoEM data processing

Movie frame alignment, estimation of the microscope CTF, particle picking, and extraction (with a downsampled pixel size of 1.686 Å and box size of 260 pixels^2^) were completed using WARP^41^. Reference-free 2D classification was performed using cryoSPARC to select for well-defined particle images^42^. These selected particles were then used for 3D classification with 50 iterations (angular sampling 7.5° for 25 iterations followed by 1.8° with local search for 25 iterations) using Relion and a previously reported closed model for the SARS-CoV-2 spike ectodomain (PBD: 6VXX) as the initial model without imposing any symmetry. 3D refinements were carried out using non-uniform refinement along with per-particle defocus refinement in cryoSPARC^43^ after which particles images were subjected to Bayesian polishing using Relion^44^ and re-extracted with a box size of 512 pixels and a pixel size of 1 Å. Another round of non-uniform refinement followed by per-particle defocus refinement followed by another non-uniform refinement was conducted in cryoSPARC. Next, 86 optics groups were defined based on the beamtilt angle used for data collection and another round of non-uniform refinement with global and per-particle defocus refinement concurrently was conducted in cryoSPARC. To better resolve the RBD, focus 3D classification was carried out using symmetry expanded particles and a mask over residues 440-452 and 495-505 of the RBD using a tau factor of 200 in Relion^45,46^. Particles from the classes with the best resolved local density were selected and then subjected to local refinement using cryoSPARC. Reported resolutions are based on the gold-standard Fourier shell correlation of 0.143 criterion and Fourier shell correlation curves were corrected for the effects of soft masking by high-resolution noise substitution^47,48^.

### Model building and refinement

USCF Chimera^49^ and Coot^50^ were used to fit atomic models of the SARS-CoV-2 RBD and ectodomain (PBD: 6M0J, 7LXY). Models were refined and rebuilt into the map using Coot^50^ and Rosetta^51,52^.

### Biolayer interferometry

Biotinylated wildtype, B.1.1.7, or E406W RBD at a concentration of 5 ng/µL in 10X kinetics buffer was loaded at 30C onto pre-hydrated streptavidin biosensor to a 1 nm total shift. The loaded tips were then dipped into a 1:3 dilution series of monomeric hACE2 beginning at 900 nM, 300 nM, or 7,500 nM for 300 seconds followed by dissociation in 10X kinetics buffer for 300 seconds. The resulting data were baseline subtracted and curves were fitted using Octet Data Analysis HT software v12.0 and plotted in GraphPad Prism 9.

### Pseudotyped VSV production

E406W and wildtype pseudotyped VSV particles were produced as previously described^23,24^. Briefly, 5 × 10^6^ HEK-293T cells were seeded in 10 cm^2^ poly-D-lysine coated plates and grown overnight until they reached ∼70% confluency. The cells were then washed 5 times with Opti-MEM (Life Technologies) and transfected with 24 µg of plasmid encoding either the wildtype or E406W SARS-CoV-2 spike protein using Lipofectamine 2000 (Life Technologies). Four hours at transfection, an equal volume of DMEM supplemented with 20% FBS and 2% PenStrep was added to the cells. Twenty to 24 hours following transfection, the cells were washed 5 times with DMEM and infected with VSVΔG/Fluc. Two hours after infection, the cells were washed 5 times with DMEM and grown in DMEM supplemented with 10% FBS and 1% PenStrep along with an anti-VSV-G antibody (I1-mouse hybridoma supernatant diluted 1:25, from CRL-2700, ATCC). Twenty to 24 hours later, the supernatant was collected, clarified by centrifugation at 2,500xg for 10 minutes, filtered through a 0.45 µm filter, and concentrated 10x using a 30 kDa filter (Amicon). The resulting pseudovirus was frozen at -80°C until use.

### Sera

Blood samples were collected from individuals 7-30 days after receiving the second dose of either Pfizer’s BNT162b2 or Moderna’s mRNA-1273 COVID-19 vaccine. All study participants were enrolled in the UWARN: COVID-19 in WA study at the University of Washington. The study protocol was approved by the University of Washington Human Subjects Division Institutional Review Board (STUDY00010350).

### Neutralization assays with vaccine-elicited sera and monoclonal antibodies

For neutralization assays using vaccine-elicited sera, HEK-ACE2 cells were seeded in 96-well poly-D-lysine coated plates at a density of 30,000 cells per well and grown overnight until they reached approximately 80% confluency. E406W and wildtype pseudoviruses were diluted 1:25 in DMEM and incubated with vaccine-elicited sera for 30 minutes at room temperature. Growth media was removed from the HEK-ACE2 cells and the virus-sera mixture was added to the cells. Two hours after infection, an equal volume of DMEM supplemented with 20% and 2% PenStrep was added to each well and the cells were incubated overnight. After 20-24 hours, ONE-Glo EX (Promega) was added to each well and the cells were incubated for 5 minutes at 37°C. Luminescence values were measured using a BioTek plate reader.

For neutralization assays using monoclonal antibodies, Vero-TMPRSS2 cells were seeded in 96-well plates at a density of 18,000 cells per overnight until they reached approximately 80% confluency. Neutralizations were conducted as described above with one modification: prior to the addition of the virus-antibody mixture, Vero-TMPRSS2 cells were washed 3 times with DMEM.

Luminescence readings from the neutralization assays were normalized and analyzed using GraphPad Prism 9. The relative light unit (RLU) values recorded from uninfected cells were used to define 0% infectivity and RLU values recorded from cells infected with pseudovirus without sera or antibodies were used to define 100% infectivity. ID50 and IC50 values for sera and monoclonal antibodies, respectively were determined from the normalized data points using a [inhibitor] vs. normalized response – variable slope model.

---

**Figure S1.**
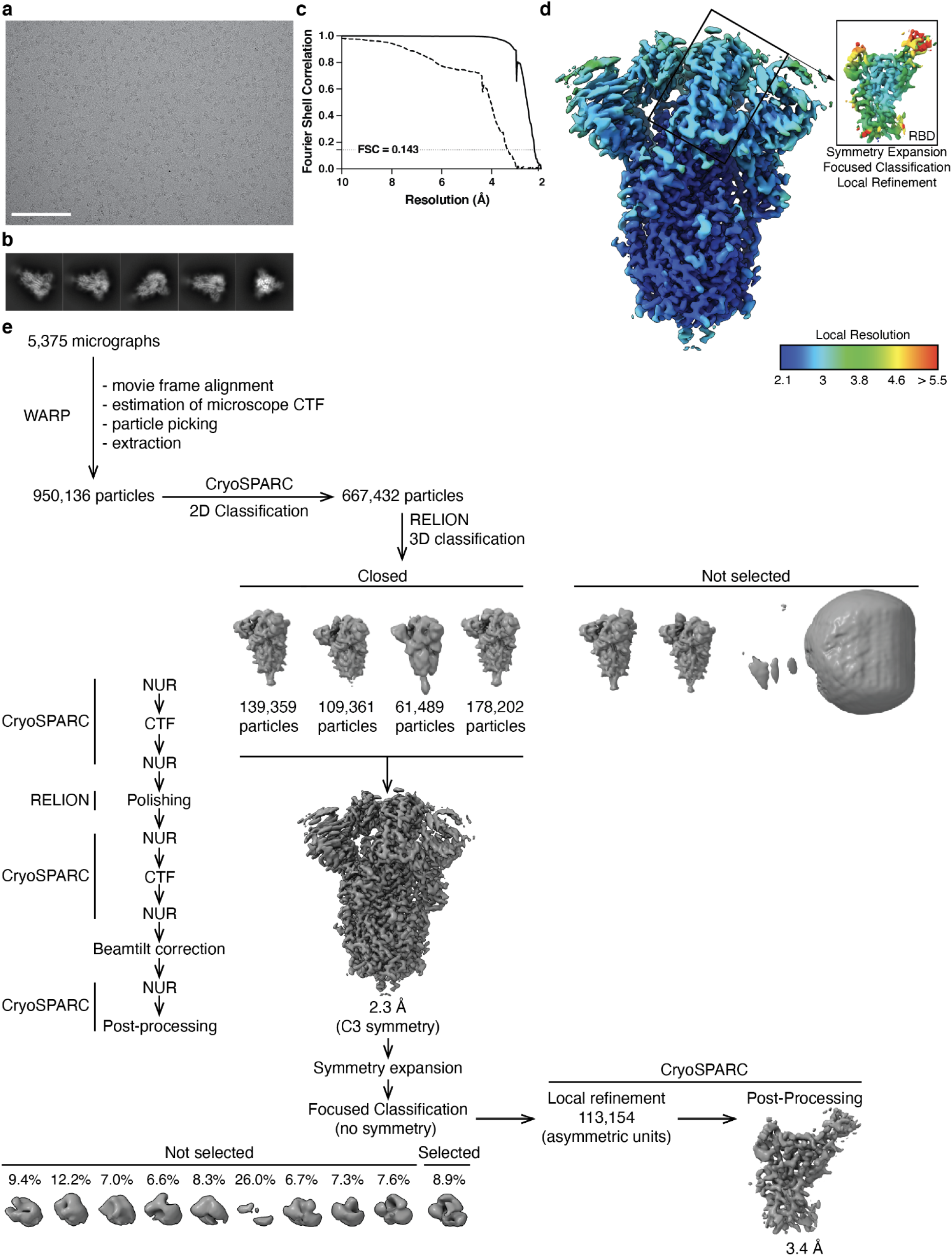
CryoEM processing and validation for the SARS-CoV-2 E406W spike dataset. **a-b**, Representative electron micrograph (a) and 2D class averages (b) obtained for the SARS-CoV-2 E406W spike ectodomain. Scale bar: 100 nm. (c) Gold-standard fourier shell correlation curves for the closed E406W S trimer (solid line) and locally refined E406W RBD (dashed line). **d**, Local resolution calculated using CryoSPARC for the E406W S ectodomain trimer (left, unsharpened map) and the locally refined RBD (right, sharpened map). **e**, CryoEM processing workflow.

**Figure S2.**
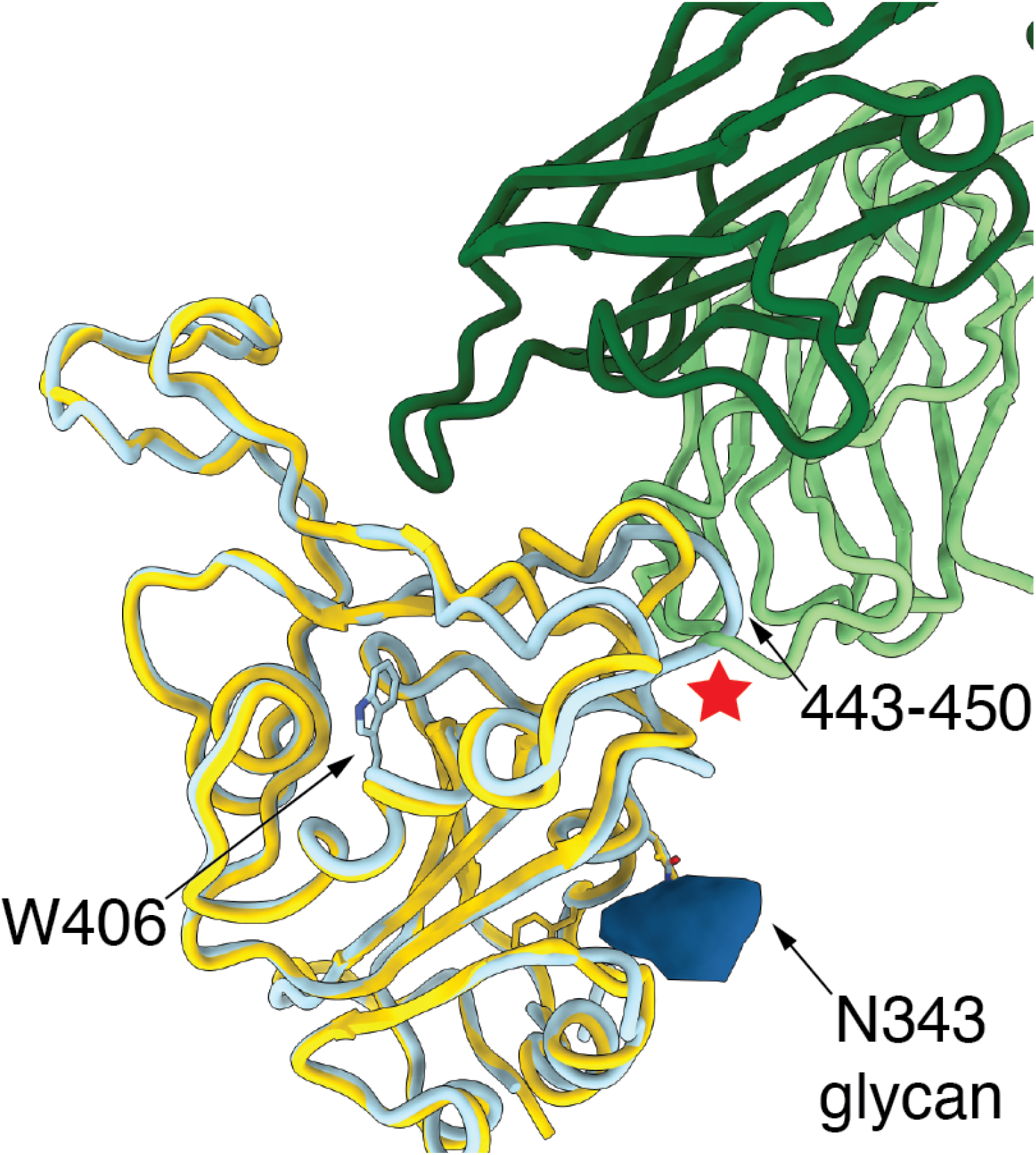
Repositioning of residues 444-450 in the W406 RBD interferes sterically with cilgavimab binding. Structural superimposition of the cilgavimab (AZD1061)-bound Wuhan-Hu-1 RBD (E406, gold, PBD 7L7E) and the W406 RBD (light blue). Key reorganized regions are labeled and the steric clash is indicated by a red star.

**Figure S3.**
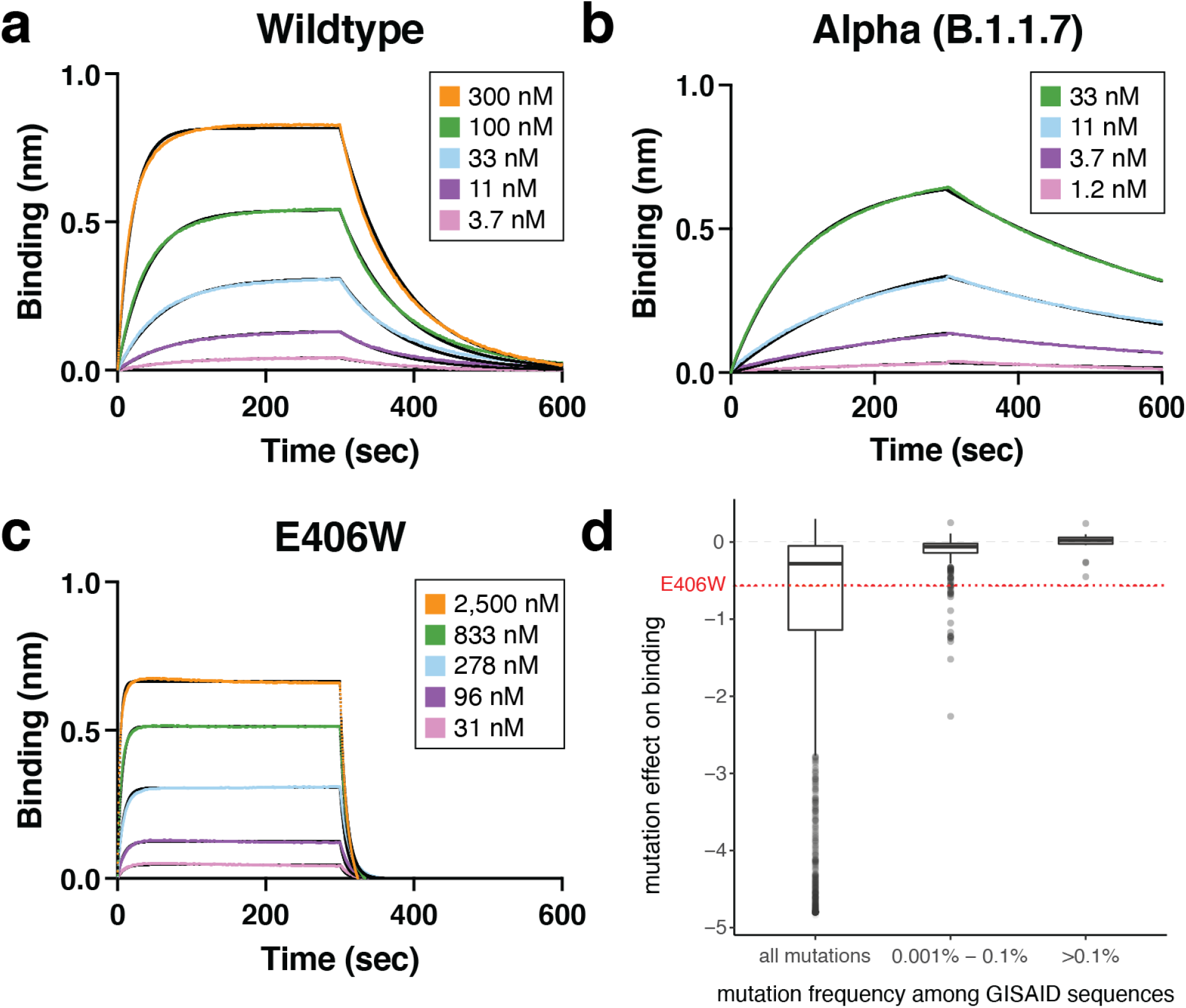
The E406W mutation dampens ACE2 binding severely. **a-c** Biolayer interferometry binding analysis of monomeric human ACE2 to immobilized Wuhan-Hu-1 (a), Alpha (N501Y, b), or E406W (c) RBDs. **d** Mutation effects on avidity for dimeric human ACE2 as measured by yeast surface display^17^ for the E406W mutation and RBD mutations found in human-derived SARS-CoV-2 isolates deposited in GISAID as of 27 September 2021 across increasing frequency thresholds.

**Figure S4.**
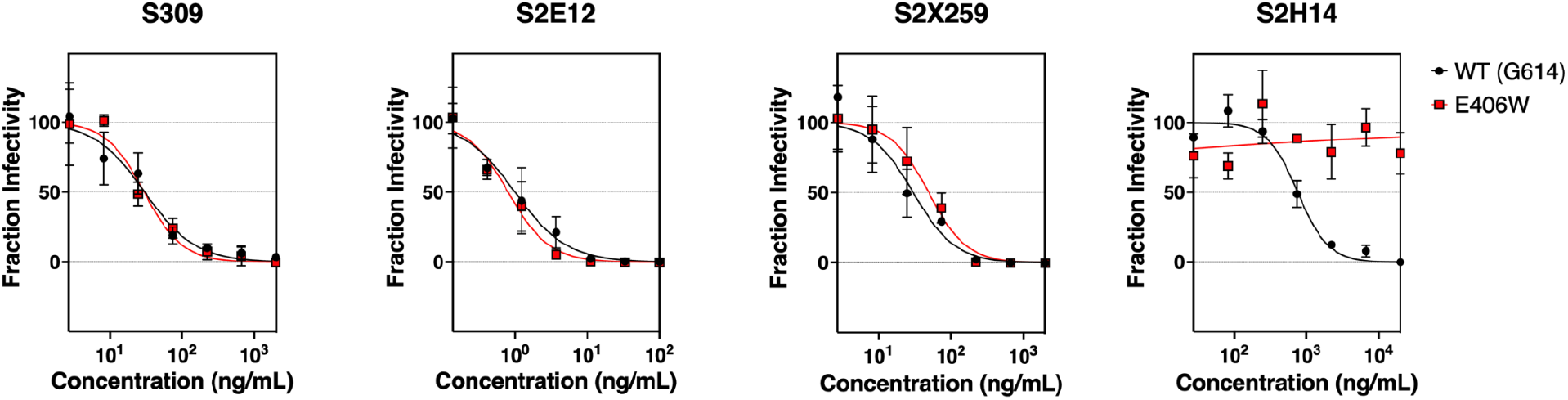
Neutralization curves for E406W/G614, shown in red, or wildtype (G164), shown in black, pseudotyped VSV using four monoclonal antibodies targeting the SARS-CoV-2 RBD. Neutralization assays were performed in triplicate and replicated twice with two batches of pseudovirus.

**Figure S5.**
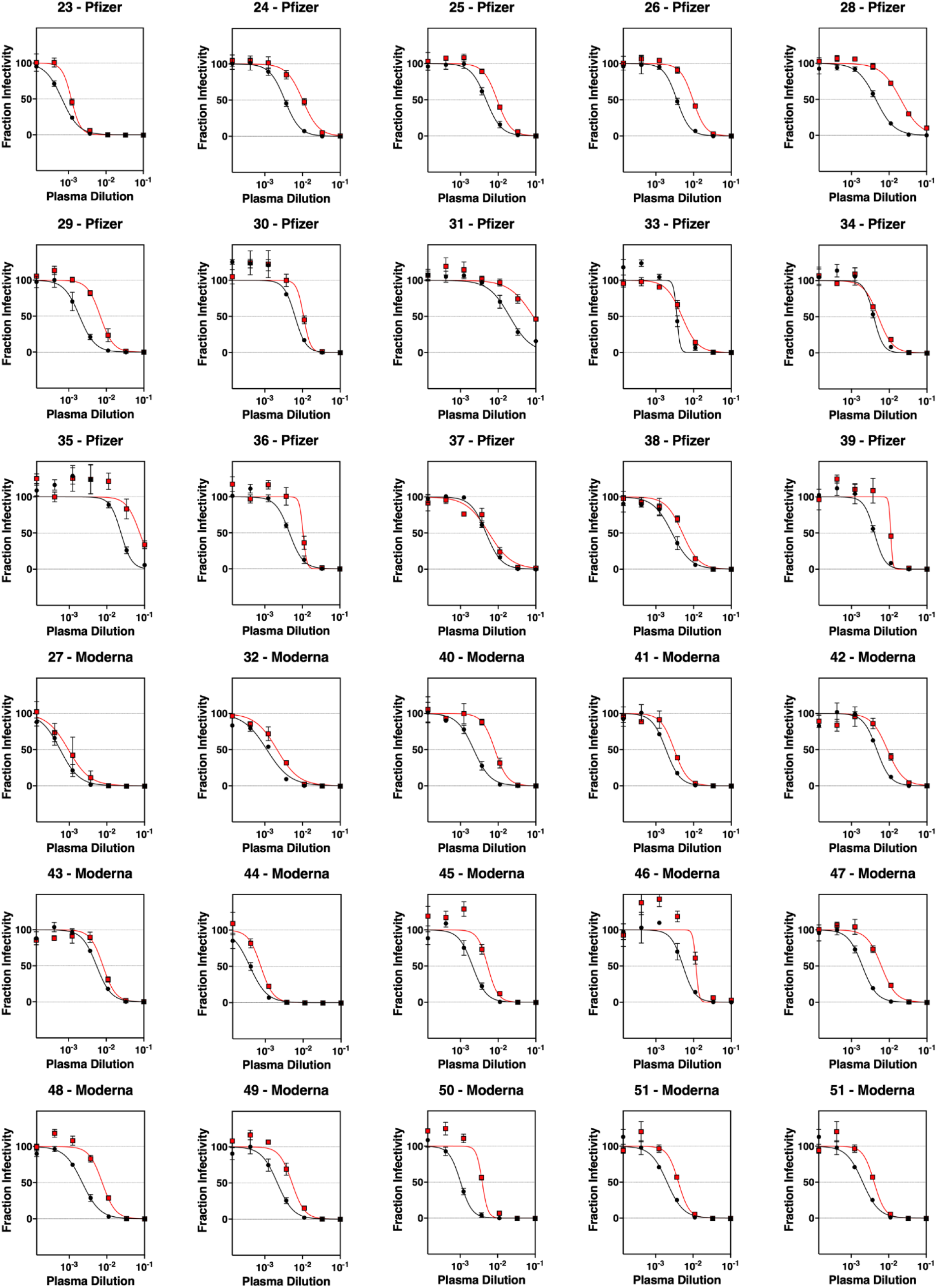
Neutralization curves against E406W/G614 S, shown in red, or wildtype (G614) S, shown in black, pseudotyped VSV for 30 sera samples collected from individuals vaccinated with either Pfizers BNT162b2 or Moderna mRNA-1273 COVID-19 vaccines. Neutralization assays were performed in triplicate and repeated at least twice with at least two distinct batches of pseudovirus.

**Table S1.** Cryo-EM data collection, refinement and validation statistics.

|  | SARS-CoV-2 S<br>E406W Ectodomain<br>(EMDB-xxxx)<br>(PDB xxxx) | SARS-CoV-2 S<br>E406W RBD<br>(local<br>refinement)<br>(EMDB-xxxx)<br>(PDB xxxx) |
| --- | --- | --- |
| Data collection and processing |  |  |
| Magnification | 105,000 | 105,000 |
| Voltage (kV) | 300 | 300 |
| Electron exposure (e-/Å<sup>2</sup>) | 63 | 63 |
| Defocus range (µm) | 0-2.6 | 0-2.6 |
| Pixel size (Å) | 0.843 | 0.843 |
| Symmetry imposed | C3 | C1 |
| Initial particle images (no.) | 950,136 | 1,281,585 |
| Final particle images (no.) | 427,195 | 113,154 |
| Map resolution (Å) | 2.3 | 3.4 |
| FSC threshold | 0.143 | 0.143 |
| Map resolution range (Å) | 2.2-8.9 | 2.8-9.4 |
| Refinement |  |  |
| Initial model used (PDB code) | 7LXY | 6M0J |
| Model resolution (Å) | 2.4 | 3.5 |
| FSC threshold | 0.143 | 0.143 |
| Map sharpening *B* factor (Å<sup>2</sup>) | -72 | -83 |
| Model composition |  |  |
| Non-hydrogen atoms | 24,198 | 1,555 |
| Protein residues | 2,994 | 194 |
| Ligands | 54 | 1 |
| *B* factors (Å<sup>2</sup>) |  |  |
| Protein | 21.13 | 41.62 |
| Ligand | 19.14 | 30 |
| R.m.s. deviations |  |  |
| Bond lengths (Å) | 0.014 | 0.013 |
| Bond angles (°) | 1.450 | 2.065 |
| Validation |  |  |
| MolProbity score | 1.36 | 1.73 |
| Clashscore | 3.94 | 4.30 |
| Poor rotamers (%) | 0 | 0 |
| Ramachandran plot |  |  |
| Favored (%) | 96.95 | 91.15 |
| Allowed (%) | 3.05 | 8.85 |
| Disallowed (%) | 0 | 0 |

**Table S2.**
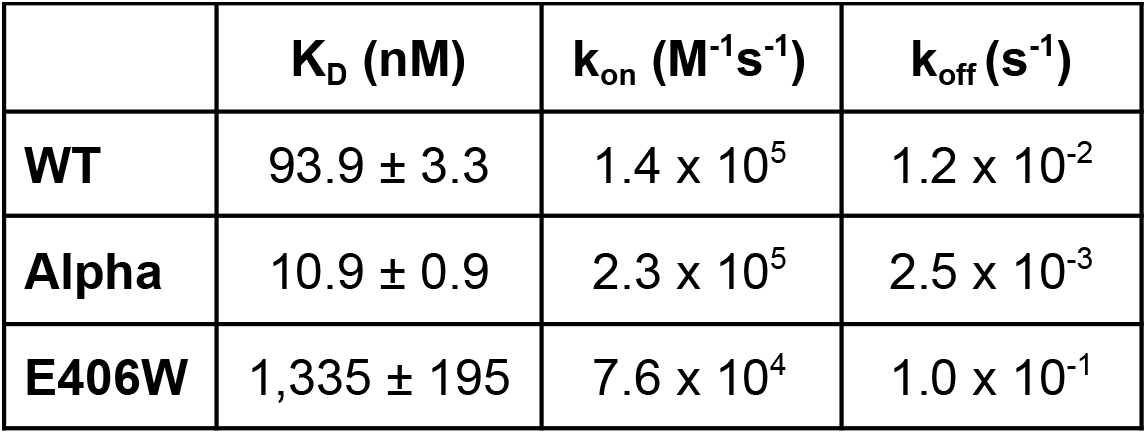
Binding kinetics of the RBD to human ACE2 as measured by biolayer interferometry. Values are presented as mean ± standard error.

**Table S3.** IC50 values for the four monoclonal antibodies tested against wildtype (G164) and E406W pseudoviruses. Values are presented as mean ± standard error.

|  | **IC<sub>50</sub> against WT pseudovirus (ng/mL)** | **IC<sub>50</sub> against E406W pseudovirus (ng/mL)** |
| --- | --- | --- |
| **S309** | 26.6 ± 3.7 | 27.3 ± 2.4 |
| **S2E12** | 0.81 ± 0.19 | 0.98 ± 0.17 |
| **S2X259** | 39.4 ± 9.2 | 33.8 ± 15.6 |
| **S2H14** | 535 ± 224 | > 20,000 |

**Table S4.** Demographic information for vaccine-elicited sera donors.

| **Study ID** | **Age** | **Vaccine Type** | **Days after second vaccination** | **Sex** | **Race** | **Ethnicity** |
| --- | --- | --- | --- | --- | --- | --- |
| 23 | 60 | Pfizer | 11 | M | White | Not Hispanic or Latino |
| 24 | 65 | Pfizer | 10 | M | White | Not Hispanic or Latino |
| 25 | 55 | Pfizer | 18 | M | White | Not Hispanic or Latino |
| 26 | 42 | Pfizer | 9 | F | White | Not Hispanic or Latino |
| 27 | 66 | Moderna | 8 | F | White | Not Hispanic or Latino |
| 28 | 63 | Pfizer | 10 | M | White | Not Hispanic or Latino |
| 29 | 27 | Pfizer | 8 | F | White | Not Hispanic or Latino |
| 30 | 38 | Pfizer | 8 | F | Asian | Not Hispanic or Latino |
| 31 | 37 | Pfizer | 21 | F | Black | Not Hispanic or Latino |
| 32 | 36 | Moderna | 7 | M | White | Not Hispanic or Latino |
| 33 | 62 | Pfizer | 15 | M | Pacific Islander | Not Hispanic or Latino |
| 34 | 54 | Pfizer | 14 | F | White | Not Hispanic or Latino |
| 35 | 60 | Pfizer | 14 | F | White | Not Hispanic or Latino |
| 36 | 32 | Pfizer | 13 | F | White | Not Hispanic or Latino |
| 37 | 52 | Pfizer | 11 | M | White | Not Hispanic or Latino |
| 38 | 61 | Pfizer | 9 | M | White | Not Hispanic or Latino |
| 39 | 32 | Pfizer | 22 | F | White | Not Hispanic or Latino |
| 40 | 40 | Moderna | 20 | M | White | Not Hispanic or Latino |
| 41 | 64 | Moderna | 16 | M | White | Not Hispanic or Latino |
| 42 | 34 | Moderna | 23 | F | Asian | Not Hispanic or Latino |
| 43 | 22 | Moderna | 20 | F | White | Not Hispanic or Latino |
| 44 | 24 | Moderna | 18 | F | White | Not Hispanic or Latino |
| 45 | 35 | Moderna | 20 | M | White | Not Hispanic or Latino |
| 46 | 40 | Moderna | 24 | M | White | Not Hispanic or Latino |
| 47 | 55 | Moderna | 20 | M | White | Not Hispanic or Latino |
| 48 | 25 | Moderna | 22 | M | White and Asian | Not Hispanic or Latino |
| 49 | 26 | Moderna | 18 | F | White | Not Hispanic or Latino |
| 50 | 36 | Moderna | 27 | F | Asian | Not Hispanic or Latino |
| 51 | 53 | Moderna | 20 | F | White | Not Hispanic or Latino |
| 52 | 47 | Moderna | 21 | M | White | Not Hispanic or Latino |

## References

1. Walls, A. C. et al. Structure, Function, and Antigenicity of the SARS-CoV-2 Spike Glycoprotein. Cell 181, 281–292.e6 (2020).

2. Zhou, P. et al. A pneumonia outbreak associated with a new coronavirus of probable bat origin. Nature (2020) doi:10.1038/s41586-020-2012-7.

3. Hoffmann, M. et al. SARS-CoV-2 Cell Entry Depends on ACE2 and TMPRSS2 and Is Blocked by a Clinically Proven Protease Inhibitor. Cell 181, 271–280.e8 (2020).

4. Corti, D., Purcell, L. A., Snell, G. & Veesler, D. Tackling COVID-19 with neutralizing monoclonal antibodies. Cell (2021) doi:10.1016/j.cell.2021.05.005.

5. Piccoli, L. et al. Mapping Neutralizing and Immunodominant Sites on the SARS-CoV-2 Spike Receptor-Binding Domain by Structure-Guided High-Resolution Serology. Cell 183, 1024–1042.e21 (2020).

6. Greaney, A. J. et al. Antibodies elicited by mRNA-1273 vaccination bind more broadly to the receptor binding domain than do those from SARS-CoV-2 infection. Sci. Transl. Med. 13, (2021).

7. McCallum, M. et al. N-terminal domain antigenic mapping reveals a site of vulnerability for SARS-CoV-2. Cell (2021) doi:10.1016/j.cell.2021.03.028.

8. Stamatatos, L. et al. mRNA vaccination boosts cross-variant neutralizing antibodies elicited by SARS-CoV-2 infection. Science (2021) doi:10.1126/science.abg9175.

9. Starr, T. N. et al. Prospective mapping of viral mutations that escape antibodies used to treat COVID-19. Science 371, 850–854 (2021).

10. Starr, T. N. et al. SARS-CoV-2 RBD antibodies that maximize breadth and resistance to escape. Nature (2021) doi:10.1038/s41586-021-03807-6.

11. Hansen, J. et al. Studies in humanized mice and convalescent humans yield a SARS-CoV-2 antibody cocktail. Science (2020) doi:10.1126/science.abd0827.

12. Baum, A. et al. Antibody cocktail to SARS-CoV-2 spike protein prevents rapid mutational escape seen with individual antibodies. Science (2020) doi:10.1126/science.abd0831.

13. Dong, J. et al. Genetic and structural basis for SARS-CoV-2 variant neutralization by a two-antibody cocktail. Nat Microbiol 6, 1233–1244 (2021).

14. Lan, J. et al. Structure of the SARS-CoV-2 spike receptor-binding domain bound to the ACE2 receptor. Nature (2020) doi:10.1038/s41586-020-2180-5.

15. Wrobel, A. G. et al. SARS-CoV-2 and bat RaTG13 spike glycoprotein structures inform on virus evolution and furin-cleavage effects. Nat. Struct. Mol. Biol. 27, 763–767 (2020).

16. Park, Y.-J. et al. Antibody-mediated broad sarbecovirus neutralization through ACE2 molecular mimicry. doi:10.1101/2021.10.13.464254.

17. Starr, T. N. et al. Deep Mutational Scanning of SARS-CoV-2 Receptor Binding Domain Reveals Constraints on Folding and ACE2 Binding. Cell 182, 1295–1310.e20 (2020).

18. Tortorici, M. A. et al. Broad sarbecovirus neutralization by a human monoclonal antibody. Nature (2021) doi:10.1038/s41586-021-03817-4.

19. Pinto, D. et al. Cross-neutralization of SARS-CoV-2 by a human monoclonal SARS-CoV antibody. Nature 583, 290–295 (2020).

20. Tortorici, M. A. et al. Ultrapotent human antibodies protect against SARS-CoV-2 challenge via multiple mechanisms. Science 370, 950–957 (2020).

21. Jette, C. A. et al. Broad cross-reactivity across sarbecoviruses exhibited by a subset of COVID-19 donor-derived neutralizing antibodies. Cell Rep. 36, 109760 (2021).

22. Martinez, D. R. et al. A broadly cross-reactive antibody neutralizes and protects against sarbecovirus challenge in mice. Sci. Transl. Med. eabj7125 (2021).

23. McCallum, M. et al. SARS-CoV-2 immune evasion by the B.1.427/B.1.429 variant of concern. Science (2021) doi:10.1126/science.abi7994.

24. McCallum, M. et al. Molecular basis of immune evasion by the Delta and Kappa SARS-CoV-2 variants. Science eabl8506 (2021).

25. Davies, N. G. et al. Estimated transmissibility and impact of SARS-CoV-2 lineage B.1.1.7 in England. Science (2021) doi:10.1126/science.abg3055.

26. Tegally, H. et al. Emergence of a SARS-CoV-2 variant of concern with mutations in spike glycoprotein. Nature (2021) doi:10.1038/s41586-021-03402-9.

27. Deng, X. et al. Transmission, infectivity, and neutralization of a spike L452R SARS-CoV-2 variant. Cell (2021) doi:10.1016/j.cell.2021.04.025.

28. Faria, N. R. et al. Genomics and epidemiology of the P.1 SARS-CoV-2 lineage in Manaus, Brazil. Science 372, 815–821 (2021).

29. Thomson, E. C. et al. Circulating SARS-CoV-2 spike N439K variants maintain fitness while evading antibody-mediated immunity. Cell (2021) doi:10.1016/j.cell.2021.01.037.

30. Collier, D. A. et al. Sensitivity of SARS-CoV-2 B.1.1.7 to mRNA vaccine-elicited antibodies. Nature (2021) doi:10.1038/s41586-021-03412-7.

31. Cele, S. et al. Escape of SARS-CoV-2 501Y.V2 from neutralization by convalescent plasma. Nature (2021) doi:10.1038/s41586-021-03471-w.

32. Wibmer, C. K. et al. SARS-CoV-2 501Y.V2 escapes neutralization by South African COVID-19 donor plasma. Nat. Med. (2021) doi:10.1038/s41591-021-01285-x.

33. Edara, V.-V. et al. Infection and Vaccine-Induced Neutralizing-Antibody Responses to the SARS-CoV-2 B.1.617 Variants. N. Engl. J. Med. (2021) doi:10.1056/NEJMc2107799.

34. Liu, C. et al. Reduced neutralization of SARS-CoV-2 B.1.617 by vaccine and convalescent serum. Cell (2021) doi:10.1016/j.cell.2021.06.020.

35. Plante, J. A. et al. Spike mutation D614G alters SARS-CoV-2 fitness. Nature (2020) doi:10.1038/s41586-020-2895-3.

36. Liu, Y. et al. Delta spike P681R mutation enhances SARS-CoV-2 fitness over Alpha variant. bioRxiv (2021) doi:10.1101/2021.08.12.456173.

37. Saito, A. et al. Enhanced fusogenicity and pathogenicity of SARS-CoV-2 Delta P681R mutation. Nature (2021) doi:10.1038/s41586-021-04266-9.

38. Crawford, K. H. D. et al. Protocol and Reagents for Pseudotyping Lentiviral Particles with SARS-CoV-2 Spike Protein for Neutralization Assays. Viruses 12, (2020).

39. Hsieh, C.-L. et al. Structure-based design of prefusion-stabilized SARS-CoV-2 spikes. Science 369, 1501–1505 (2020).

40. Suloway, C. et al. Automated molecular microscopy: the new Leginon system. J. Struct. Biol. 151, 41–60 (2005).

41. Tegunov, D. & Cramer, P. Real-time cryo-electron microscopy data preprocessing with Warp. Nat. Methods 16, 1146–1152 (2019).

42. Punjani, A., Rubinstein, J. L., Fleet, D. J. & Brubaker, M. A. cryoSPARC: algorithms for rapid unsupervised cryo-EM structure determination. Nat. Methods 14, 290–296 (2017).

43. Punjani, A., Zhang, H. & Fleet, D. J. Non-uniform refinement: adaptive regularization improves single-particle cryo-EM reconstruction. Nat. Methods 17, 1214–1221 (2020).

44. Zivanov, J., Nakane, T. & Scheres, S. H. W. A Bayesian approach to beam-induced motion correction in cryo-EM single-particle analysis. IUCrJ 6, 5–17 (2019).

45. Zivanov, J. et al. New tools for automated high-resolution cryo-EM structure determination in RELION-3. Elife 7, (2018).

46. Scheres, S. H. W. RELION: implementation of a Bayesian approach to cryo-EM structure determination. J. Struct. Biol. 180, 519–530 (2012).

47. Rosenthal, P. B. & Henderson, R. Optimal determination of particle orientation, absolute hand, and contrast loss in single-particle electron cryomicroscopy. J. Mol. Biol. 333, 721–745 (2003).

48. Chen, S. et al. High-resolution noise substitution to measure overfitting and validate resolution in 3D structure determination by single particle electron cryomicroscopy. Ultramicroscopy 135, 24–35 (2013).

49. Pettersen, E. F. et al. UCSF Chimera--a visualization system for exploratory research and analysis. J. Comput. Chem. 25, 1605–1612 (2004).

50. Emsley, P., Lohkamp, B., Scott, W. G. & Cowtan, K. Features and development of Coot. Acta Crystallogr. D Biol. Crystallogr. 66, 486–501 (2010).

51. Frenz, B. et al. Automatically Fixing Errors in Glycoprotein Structures with Rosetta. Structure 27, 134–139.e3 (2019).

52. Wang, R. Y.-R. et al. Automated structure refinement of macromolecular assemblies from cryo-EM maps using Rosetta. Elife 5, (2016).

